# Identification of known and novel long non-coding RNAs potentially responsible for the effects of BMD GWAS loci

**DOI:** 10.1101/2021.11.04.467171

**Authors:** Abdullah Abood, Larry Mesner, Will Rosenow, Basel M. Al-Barghouthi, Nina Horwitz, Elise F. Morgan, Louis C. Gerstenfeld, Charles R. Farber

## Abstract

Osteoporosis, characterized by low bone mineral density (BMD), is the most common complex disease affecting bone and constitutes a major societal health problem. Genome-wide association studies (GWASs) have identified over 1100 associations influencing BMD. It has been shown that perturbations to long non-coding RNAs (lncRNAs) influence BMD and the activities of bone cells; however, the extent to which lncRNAs are involved in the genetic regulation of BMD is unknown. Here, we combined the analysis of allelic imbalance (AI) in human acetabular bone fragments with a transcriptome-wide association study (TWAS) and expression quantitative trait loci (eQTL) colocalization analysis using data from the Genotype-Tissue Expression (GTEx) project to identify lncRNAs potentially responsible for GWAS associations. We identified 27 lncRNAs in bone that are located in proximity to a BMD GWAS association and harbor SNPs demonstrating AI. Using GTEx data we identified an additional 31 lncRNAs whose expression was associated (FDR correction<0.05) with BMD through TWAS and had a colocalizing eQTL (regional colocalization probability (RCP)>0.1). The 58 lncRNAs are located in 43 BMD associations. To further support a causal role for the identified lncRNAs, we show that 23 of the 58 lncRNAs are differentially expressed as a function of osteoblast differentiation. Our approach identifies lncRNAs that are potentially responsible for BMD GWAS associations and suggest that lncRNAs play a role in the genetics of osteoporosis.

## Introduction

Osteoporosis is characterized by low bone mineral density (BMD) and deteriorated structural integrity which leads to an increased risk of fracture ^1,2^. In the U.S. alone, 12 million individuals have been diagnosed with osteoporosis, contributing to over 2 million fractures per year ^3^. This number is expected to nearly double by 2025, resulting in approximately $26 billion in health care expenditures ^3^.

BMD is one of the strongest predictors of fracture ^4^ and is a highly heritable quantitative trait (h^2^ = 0.5-0.8) ^5–8^. As a result, the majority of genome-wide association studies (GWASs) conducted for osteoporosis have focused on BMD. The largest BMD GWAS performed to date used the UK BioBank (N∼420K) and identified 1103 associations influencing heel estimated BMD (eBMD) ^9^. One of the main challenges of BMD GWAS is that the majority (>90%) of associations implicate non-coding variants that lie in intronic or intergenic regions suggesting they have a role in gene regulation. This has made it difficult to pinpoint causal genes and highlights the need for follow-up studies ^10^. In addition, few studies have systematically evaluated non-coding transcripts as potential causal genes.

The largest and most functionally diverse family of non-coding transcripts are long non-coding RNAs (lncRNAs). LncRNAs are transcripts longer than 200 nucleotides and have no coding potential ^11^. The majority of lncRNAs share sequence features with protein-coding genes including a 3’ poly-A tail, a 5’ methyl cap, and an open reading frame ^12^. However, their expression is low and heterogenous, and they show intermediate to high tissue specificity ^13^. Aberrant expression of lncRNAs has been linked to diseases such as osteoporosis ^14^. Additionally, there is accumulating evidence suggesting their involvement in key regulatory pathways, including osteogenic differentiation ^11,15^.

Although understudied in the context of GWAS ^13^, there is increasing evidence suggesting that lncRNAs are causal for a subset of associations identified by GWAS. A recent analysis of data from the Genotype-Tissue Expression (GTEx) project identified 690 potentially causal lncRNAs underlying associations influencing risk of a wide range of diseases ^13^. Additionally, there is emerging evidence implicating lncRNAs in the genetics of BMD ^16–18^. For example, a study reported 575 differentially expressed lncRNAs between high and low BMD groups in Caucasian women, 26 of which regulate protein-coding genes that are potentially causal in BMD GWAS ^19^. Additionally, a recent BMD single nucleotide polymorphism (SNP) prioritization analysis implicated lncRNAs as potential causal mediators ^20^. Together these studies suggest that lncRNAs may play an important role in the genetic regulation of bone mass.

In recent years, a number of approaches have been developed that utilize transcriptomics data to inform GWAS, including the analysis of allelic imbalance (AI), transcriptome-wide association studies (TWASs), and expression quantitative trait loci (eQTL) colocalization ^21^. AI results from the cis-regulatory effects (i.e., local eQTL) that can be tracked using heterozygous coding SNPs. In transcriptome-wide association studies (TWASs) the genetic component of gene expression in a reference population is estimated and then imputed in a much larger population. Once gene expression is imputed, genetically regulated gene expression is associated with a disease or disease phenotype ^22^. Most genes identified by TWAS are located in GWAS associations for that disease and, as a result, TWAS can pinpoint genes likely to be causal at GWAS loci. eQTLs are genetic variants associated with changes in gene expression and can be tissue-specific or shared across multiple tissues. eQTL colocalization tests whether the change in gene expression and the change in a trait of interest are driven by the same shared genetic variant(s). All three approaches, alone or in combination, have been successfully used to pinpoint potential causal disease genes at GWAS associations.

Here, we identified lncRNAs that are potentially responsible for the effects of BMD GWAS associations by first applying AI to bone samples and, next, applying TWAS and eQTL colocalization to gene expression data from GTEx. Through both approaches we identified 58 lncRNAs with evidence of being causal BMD GWAS genes. We further prioritized these lncRNAs by identifying those that were differentially expressed as a function of osteoblast differentiation. Together, these results highlight the potential importance of lncRNAs as candidate causal BMD GWAS genes.

## Results

In this study, we used two approaches to identify lncRNAs that potentially underlie BMD GWAS associations. In the first approach, we quantified known and novel lncRNAs using RNA-seq data from human bone fragments and identified lncRNAs located in proximity of a BMD GWAS association and harboring SNPs demonstrating AI. In the second approach, we leveraged GTEx to identify lncRNAs across a large number of tissues and cell-types whose expression was significantly associated with BMD by TWAS and regulated by an eQTL which colocalized with a BMD association. **Figure 1** provides an overview of our study.

**Figure 1:**
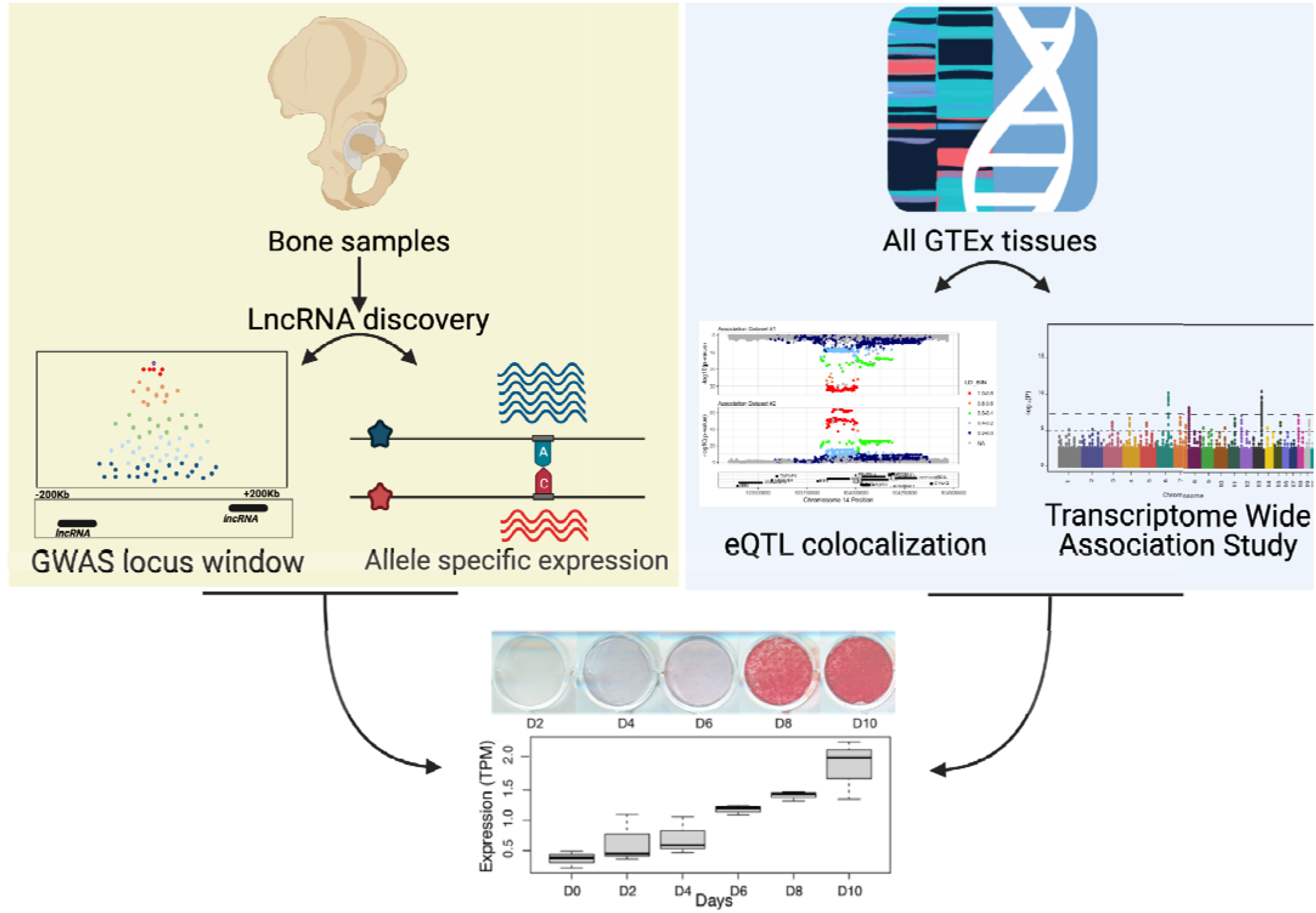
Overview of the study. We conducted de novo lncRNA discovery using RNA-seq data on human acetabular bone fragments from 17 patients. We then identified known and novel lncRNAs located in GWAS associations that were influenced by Allelic Imbalance (AI) (yellow box). We applied Transcriptome Wide Association Study (TWAS) and colocalization on eQTL data from 49 Genotype-Tissue Expression (GTEx) project tissues (blue box). We assessed the role of lncRNAs reported by both approaches in osteogenic differentiation using RNAseq data from the human fetal osteoblast (hFOB) cell line at six time points across differentiation (bottom panel).

### Generation of bone expression data from bone fragments

To identify potentially casual lncRNAs in a BMD relevant tissue, we generated total RNA-seq (ribo-depleted) data on bone fragments isolated from acetabular reamings from patients undergoing hip arthroplasty (N=17; 5 males and 12 females; ages 43 to 80). In contrast to most gene expression data generated on bone which are typically from biopsies that contain marrow, we were able to remove the marrow leaving purified trabecular and cortical bone. We hypothesized that the acetabular bone fragments consisted primarily of late-stage osteoblasts/osteocytes ^23^, allowing us to characterize lncRNAs enriched in these cell types. To confirm that the acetabular samples were enriched in osteocytes, we compared these data to published RNA-seq data on bone biopsies ^24^. Farr et al. generated RNA-seq data on 58 iliac crest needle biopsies from healthy women containing both bone and marrow. Average transcripts per million (TPM) across all samples in both experiments were highly correlated (**Figure 2A**, r=0.845, P < 2.2 × 10^−16^). Importantly, differential expression analysis between the two datasets showed that the top 1000 genes with the largest fold change increase in the bone fragment samples compared to bone biopsy samples were enriched in Gene Ontology (GO) terms such as “skeletal system development” (FDR=4.01 × 10^−3^) and “extracellular matrix organization” (FDR=4.11 × 10^−5^).

**Figure 2:**
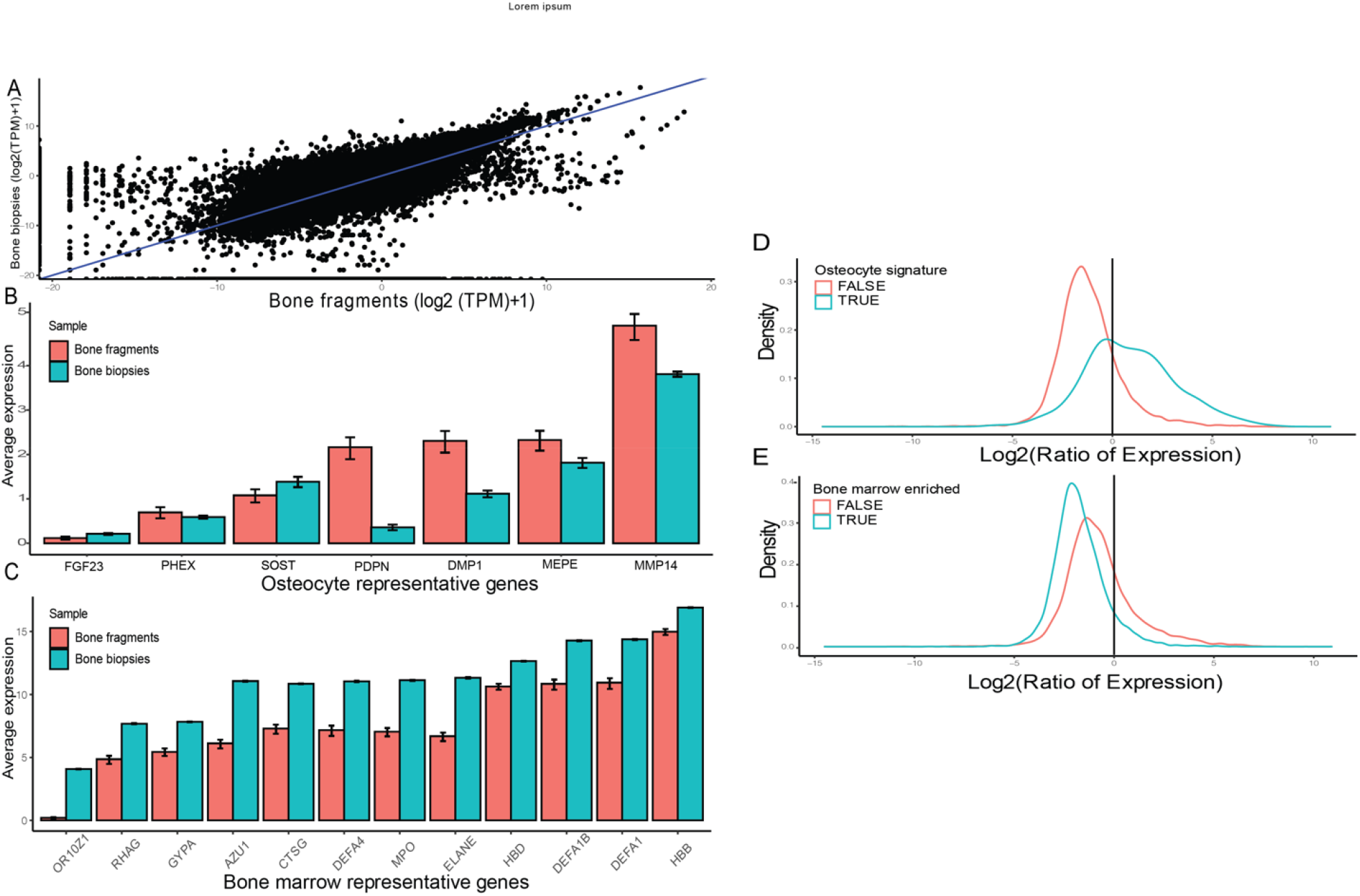
Enrichment of osteocyte marker genes in bone fragment samples (used in this study) compared to bone biopsy samples in the literature. A) Overall gene expression is highly correlated between the RNA-seq data generated in both studies (r=0.845, P < 2.2 × 10-16) 24 B) Gene expression of osteocyte marker genes reported in 23 showing enrichment in the bone fragments samples (this study) relevant to bone biopsies. C) Gene expression of bone marrow enriched genes reported in www.proteinatlas.org/ showing higher expression in bone biopsy samples. D) Osteocyte signature genes reported in Youlten et al. 25 are highly expressed in bone fragment samples relative to bone biopsies E) Bone marrow enriched genes reported in 25 are highly expressed in bone biopsy samples compared to bone fragment samples.

To support the notion that our samples are unique in osteocyte enrichment, we used data from a recent study that identified an osteocyte gene signature consisting of 1239 genes in mice and their orthologs in humans ^25^. The ratio of expression (bone fragment samples / bone biopsy samples) was used. A ratio value > 1 indicates that gene expression is higher in the bone fragment samples relative to the bone biopsy samples. In contrast, a ratio value < 1 indicates that the gene is highly expressed in bone biopsy samples relative to bone fragment samples. We expect to see that osteocyte signature genes show ratio values > 1 and marrow enriched genes show ratio values < 1. The osteocyte signature genes showed a median ratio of 1.72 (62% of osteocyte signature genes ratio > 1). Additionally, the ratio of expression of genes enriched in bone marrow showed a median of 0.27 (91% of marrow enriched genes ratio < 1). The distribution of osteocyte signature genes ratio values showed a significant median shift (Wilcoxon test, P < 2.2 × 10^−16^) (**Figure 2D**), and the opposite pattern was observed for the bone marrow enriched genes (Wilcoxon test, P < 2.2 × 10^−16^) (**Figure 2E**). These data suggest that the purified acetabular bone fragments are enriched for late osteoblasts/osteocytes compared to iliac crest biopsies.

### Identifying novel lncRNAs in purified acetabular bone fragments

Given the paucity of bone transcriptomics data in the literature, and the tissue-specific nature of lncRNA expression, we hypothesized that many bone/osteocyte specific lncRNAs would not be present in current sequence databases. Additionally, ∼50% of lncRNAs do not possess a poly-A tail modification ^26^ and most RNA-seq data is generated after poly-A selection. Therefore, in order to capture a more comprehensive profile of lncRNAs in bone, we implemented a lncRNA discovery step to identify putative “novel” lncRNA transcripts using the computational algorithm CPAT ^27^. Across the 17 bone samples we identified 6612 known lncRNAs and 2440 novel lncRNAs (Supplementary tables 1 and 2). The mean length of novel lncRNAs was 30.3 Kb and median length of 11.8 Kb. These values were comparable to the mean length of known lncRNAs expressed in the bone samples (mean = 35.4 Kb; median = 4.7 Kb).

### Identifying potentially casual lncRNAs in bone

For lncRNAs to be considered potentially causal in bone, we identified those that are both located in proximity of a BMD GWAS association and regulated by AI. We hypothesized that such genes may be causal for their respective associations because of the potential to be regulated by an eQTL which colocalizes with a BMD association. Of the 9,052 lncRNAs (2440 novel and 6612 known) we quantified in acetabular bone, 1,496 lncRNAs (∼17% of expressed lncRNAs) were found within a 400Kb window (± 200Kb from the lncRNA start site) of each of 1103 GWAS associations previously identified by Morris et al. ^9^.

Next, we identified heterozygous coding variants that demonstrated significant evidence of AI within lncRNAs. Of the total number of lncRNAs we identified, 174 (47 known, 127 novel; ∼2% of expressed lncRNAs) had at least one SNP demonstrating AI in at least one of the 17 bone fragment samples. Out of the 174, 27 (15.5%; 8 known, 19 novel) were located in proximity of a GWAS association (**Figure 3A, Supplementary Table 3**).

**Figure 3:**
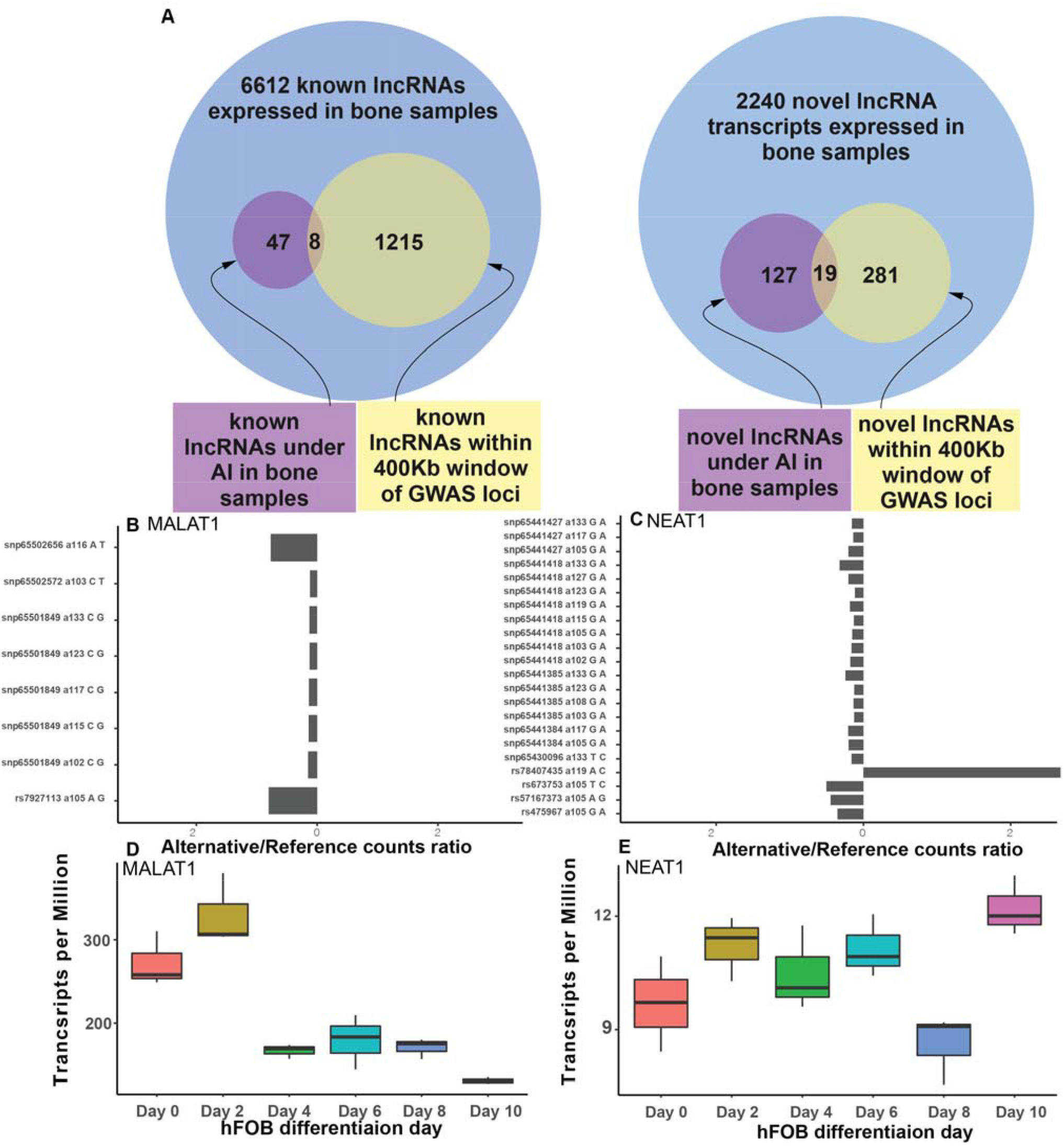
Identification of lncRNAs located within eBMD GWAS associations, are under AI in acetabular bone, and are differentially expressed in hFOBs. A) Venn diagram showing the number of known and novel lncRNAs within proximity of GWAS loci, implicated by AI, and implicated by both approaches. B) lncRNA MALAT1 AI plot showing the ratio of reads aligning to the alternative SNP relative to the reference SNP in eight of the bone fragments samples where the gene is under AI. C) lncRNA NEAT1 AI plot showing the ratio of reads aligning to the alternative SNP relative to the reference SNP in ten of the bone fragments samples where the gene is under AI. rs78407435 is not in LD with the rest of the SNPs in the region and this is likely the reason it shows a different direction of effect. D) Expression of MALAT1 across hFOB differentiation points. E) Expression of NEAT1 across hFOB differentiation points.

### Identifying putatively causal lncRNAs by leveraging GTEx

Next, we sought to leverage non-bone data to identify potentially causal lncRNAs. To do this, we integrated 1103 BMD GWAS loci ^9^ with GTEx (v8) eQTL data by coupling TWAS ^28^ using S-MultiXScan ^29^ and Bayesian colocalization analysis using fastENLOC ^30^. The rationale behind using GTEx data is genes that are shared in multiple tissues and showing a colocalizing eQTL with BMD GWAS data can be potentially causal in bone tissue as well. Our TWAS analysis resulted in 333 significant lncRNA-BMD associations (FDR correction < 0.05). Our colocalization analysis yielded 48 lncRNAs with a colocalizing eQTL (RCP > 0.1) in at least one GTEx tissue. There were 31 lncRNAs significant in both the TWAS and eQTL colocalization analysis (**Supplementary Table 4**).

### Most identified lncRNAs are the only potential causal mediators implicated by TWAS/eQTL colocalization in their respective GWAS associations

To determine if the lncRNAs listed in **Supplementary Table 4** are the strongest candidates in their respective GWAS associations, we evaluated a recent report of protein coding genes that used the same approach ^31^. Five out of the 31 lncRNAs (*LINC01116, LINC01117, SNHG15, LINC01290, LINC00665*) have a protein coding gene with a colocalizing eQTL (*HOXD8, HOXD9, MYO1G, NACAD, EMP2, ZFP14, ZFP82*) within 1 Mb of the lncRNA start site (**Supplementary Table 5**). Upon further investigation of the RCP values, some of the lncRNAs showed higher RCP than their protein coding gene counterpart. For example, *LINC01290* had a higher RCP in lung tissue (0.4992) compared to its counterpart EMP2 (0.2227). On the other hand, the same lncRNA has a lower RCP value (0.1498) than *EMP2* (0.6089) in breast and mammary gland tissue. However, for the remaining lncRNAs, this analysis provides support that the lncRNA alone is the potential causal mediator in the region as we show no evidence of protein coding colocalization within 1 Mb distance of the start site of the lncRNA.

### Many identified lncRNAs are differentially expressed as a function of osteoblast differentiation

To provide further support for the hypothesis that these lncRNAs mediate GWAS associations, we measured their expression as a function of osteoblast differentiation in hFOBs. We performed total RNA-seq at six hFOB differentiation time-points (Days 0, 2, 4, 6, 8, and 10). Of the 27 lncRNAs implicated in the analysis of AI, all eight known lncRNAs were differentially expressed (FDR<0.05). On the other hand, none of the novel lncRNAs were differentially expressed (**Supplementary Table 3**). Examples of the identified genes include *MALAT1* and *NEAT1* (**Figure 3B** and **3C**), which were differentially expressed in hFOBs and showed evidence of AI in 8 and 10 of the 17 acetabular bone samples, respectively. There were four unique SNPs in the exonic regions of *MALAT1* (**Figure 3B**) that were heterozygous in at least one of the 17 individuals (with a maximum of 8 individuals). All four SNPs showed higher expression in the alternative allele relative to the reference allele. The expression of *MALAT1* gene decreased as the cell differentiated into a mineralizing state. Additionally, there were nine unique SNPs reported in the exonic regions of NEAT1 that were heterozygous in at least one of the 17 individuals (with a maximum of 10 individuals). Of the nine, eight showed higher expression associated with the alternative allele compared to the reference allele. The remaining SNP was associated with the opposite pattern and this was likely due to it being the only SNP not in high LD with the others (R^2^ = 0.0021). NEAT1 showed significant increase in expression around day 10 in hFOBs.

We assessed the expression of lncRNAs identified by GTEx TWAS/eQTL coloclaization in osteoblast differentiation using the same approach in the previous section. Out of the 31 lncRNAs identified by TWAS/eQTL colocalization, 15 were found to be differentially expressed (*LINC00184, SH3RF3-AS1, LINC01116, LINC01934, C3orf35, LINC01018, ARRDC3-AS1, LINC00472, SNHG15, GAS1RR, LINC00840, LINC01537, LINC00346, LINC01415, MIR155HG*). In general, the expression of those genes in hFOBs was low compared to the lncRNAs reported in the AI section. Examples include *SHR3F3-AS1* and *LINC00472*, which were regulated by colocalizing eQTL (**Figure 4 B and D**) and were differentially expressed in hFOBs. (**Figure 4 C and E**). *SH3RF3-AS1* was shown to have the highest RCP value overall (RCP= 0.72) and in only one GTEx tissue (cultured fibroblasts) (**Figures 4A and 4D, Table 2**). While the gene was differentially expressed across hFOB differentiation points, it had a very low overall level of expression (**Figure 4E**). The pattern of expression decreased during mid differentiation points with spikes in early and late points (**Figure 4E**). *LINC00472* was shown to have a colocalizing eQTL in four GTEx tissues with the highest RCP value in brain cerebellar hemisphere (RCP = 0.37) (**Figures 4A and 4B, Table 2**). The gene also showed a moderate level of expression in hFOBs with an average of 1.5 TPM (**Figure 4C**). The expression of LINC00472 peaked at day 2 and then declined (**Figure 4C**).

**Figure 4:**
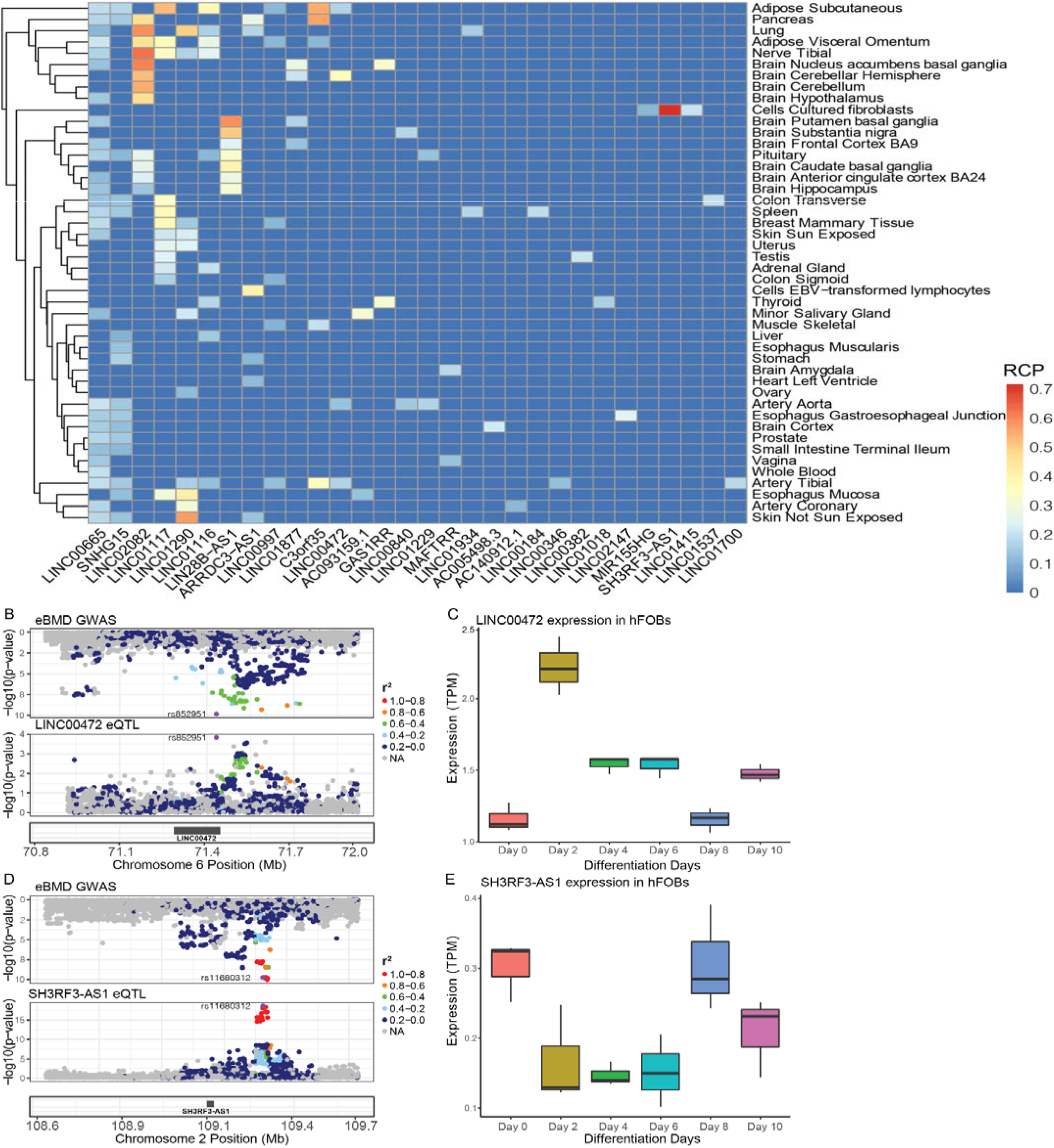
lncRNAs implicated by eQTL colocalization and TWAS are potential causal mediators of BMD GWAS loci. A) Heatmap showing colocalization events in GTEx tissues. B) lncRNA LINC00472 colocalization plot showing colocalization of eBMD GWAS locus with eQTL from Brain Cerebellar Hemisphere with RCP of 0.37 C) Differential expression of LINC00472 across hFOB differentiation points D) lncRNA SH3RF3-AS1 colocalization plot showing colocalization of eBMD GWAS locus with GTEx fibroslats eQTL data with RCP of 0.72 E) Differential expression of SH3RF3-AS1 across hFOB differentiation points.

## Discussion

In this study, we interrogated BMD GWAS loci and identified known and novel lncRNAs as potential causal mediators. We identified potentially important lncRNA using two different approaches. First, we identified novel and known lncRNAs in a unique transcriptomic bone dataset that were localized in GWAS loci and demonstrated AI. Second, we implicated additional lncRNAs by leveraging GTEx and identifying eQTLs in non-bone tissues that colocalized with eBMD GWAS loci whose expression was associated with eBMD via TWAS. We also assessed differential expression across the time course of hFOB differentiation to provide more evidence of a potential causal role for these lncRNAs.

In the first approach, we set out to perform transcriptomics on a unique sets of bone samples in order to identify novel lncRNAs in bone, provide deeper coverage for known lncRNA identification, and apply AI analysis. The bone samples that exist in the literature are from bone biopsies, and as we show in the results section, they are less enriched in bone-relevant genes compared to the dataset produced by the bone fragments used in this study.

A total of eight lncRNAs (*NEAT1, MALAT1, DLEU2, LINC01578, CARMN, AC011603*.*3, PXN-AS1, AC020656*.*1*) were found to be within a 400 Kb window of an eBMD GWAS locus and were also differentially expressed across hFOB differentiation time points. Many of these lncRNAs have been demonstrated to play a role in bone. For example, *NEAT1* has been reported to stimulate osteoclastogenesis via sponging miR-7 ^32^ and NEAT1/miR-29b-3p/BMP1 axis promotes osteogenic differentiation in human bone marrow-derived mesenchymal stem cells ^33^. In addition, *MALAT1* has been shown to influence BMD ^34^. *MALAT1* acts as a sponge of miR-34c to promote the expression of *SATB2. SATB2* then acting to reduce the ALP activity of osteoblasts and mineralized nodules formation ^34^. A recent study has shown that *LINC01578* (referred to as *CHASERR* in this study) represses chromodomain Helicase DNA Binding Protein 2 (*Chd2)*. A model for *Chd2* loss of function by the International Mouse Phenotyping Consortium (IMPC) ^35^ reported that these mice exhibit significant decreased body weight and length, skeletal abnormalities, abnormal bone structure, decreased fat levels and bone mineral density ^36^. Lastly, *DLEU2* expression has been shown to be inversely correlated with BMD in a study involving postmenopausal Caucasian women ^37^. The remaining four lncRNAs have not been reported to date to have a role in bone and should be further pursued.

In our second analysis, we reported 15 lncRNAs implicated jointly by colocalization, TWAS, and differential expression analysis. We show one example of the 15 lncRNAs reported *SH3RF3-AS1* in **(Figure 4A)**. Most of these lncRNAs have not been shown previously in the literature to have a role in bone biology. However, *LINC00472* **(Figure 4B)** has been experimentally shown to influence osteogenic differentiation by sponging miR-300 which in turn increases the expression of *Fgfr2* in mice ^38^. These preliminary results provide more evidence to the potential causal role of these lncRNAs in osteoporosis.

This study is not meant to be comprehensive as we are limited by the number of samples and are not suitably powered to identify eQTLs and apply TWAS/colocalization analysis. However, due to the scarcity of population-level bone transcriptomic dataset, and the lack of bone cell or tissue data in GTEx, our study is an attempt to systematically leverage the available datasets to capture a subset of lncRNAs that we think are potentially causal. As mentioned, some of these lncRNAs have been implicated experimentally outside of this study. Moreover, lncRNAs under AI and within proximity of GWAS loci may not be causal as they could be false positives because they are not prioritized via a systems analysis like colocalization. Another limitation of our study is that we evaluated their expression as a function of osteoblast differentiation; however, it is likely that some of the lncRNAs, if truly causal, impact BMD via a function in other cell-types (e.g., osteoclasts). Future studies should focus on enhancing these results by generating transcriptomic and eQTL datasets from bone and other bone cell types, using network approaches to aid in the prioritization of lncRNAs, and experimentally validating the role of specific lncRNAs.

In this study, we were able to use multiple systems genetics approaches on two transcriptomic datasets (acetabular bone and GTEx) to identify lncRNAs that are potentially responsible for the effects of some BMD GWAS loci. This is the first study to our knowledge that evaluated the role of lncRNAs in mediating the effect of BMD GWAS loci from a genome-wide perspective. We combined osteoblast differentiation samples and the literature to provide experimental evidence in previous studies to support the causal mediator list we generated from our analysis. These results highlight the importance of studying other aspects of the transcriptome to identify potential drug targets for osteoporosis and bone fragility.

## Data availability statement

Analysis code is available on GitHub [https://github.com/aa9gj/lncRNA_publication]. Raw samples are submitted to Gene Expression Omnibus [https://www.ncbi.nlm.nih.gov/geo/] reference number [GSE186922].

## Acknowledgements

Research reported in this publication was supported in part by the National Institute of Arthritis and Musculoskeletal and Skin Diseases of the National Institutes of Health under Award Number AR071657 to Charles R. Farber, Louis C. Gerstenfeld, and Elise F. Morgan, and Abdullah Abood was supported in part by a National Institutes of Health, Biomedical Data Sciences Training Grant (5T32LM012416). The authors acknowledge Dr. Emily Farber for generating RNAseq data on bone fragments. We thank the IMPC for accessibility to BMD data in knockout mice (www.mousephenotype.org). The Genotype-Tissue Expression (GTEx) Project was supported by the Common Fund of the Office of the Director of the National Institutes of Health, and by NCI, NHGRI, NHLBI, NIDA, NIMH, and NINDS. The data used for the analyses described in this manuscript were obtained from the GTEx Portal on 6/30/20.

## Methods

### Patient demographics

All human specimen collection was performed in accordance with IRB approval from our institution (IRB number H-32517). Acetabular reaming from 17 Boston Medical Center (BMC) patients (ages 43-80 year) undergoing elective hip arthroplasty were collected: 12 Females and 5 Males; 8 Black, 8 White, and 1 Hispanic. This demographic mix reflects the population serviced by BUMC, which is an urban safety-net hospital.

### RNA extraction

Bone fragments were isolated from the 17 patients. Total RNA was isolated from bone fragments as previously described in ^39^. Total RNA-Seq libraries were constructed from bone as well as hFOB RNA samples using Illumina TruSeq Stranded Total RNA with Ribo-Zero Gold sample prep kits. Constructed libraries contained all RNAs greater than 100 nt (both unpolyadenylated and polyadenylated) minus cytoplasmic and mitochondrial rRNAs. Samples were sequenced to achieve a minimum of 50 million reads 2 × 75 bp paired-end reads on an Illumina NextSeq500.

### Human fetal osteoblast (hFOB) cell line culture

hFOB 1.19 cells (ATCC #CRL-11372) were cultured at 34C and differentiated at 39.5C as recommended with the following modifications. Growth media: Minimal Essential Media (MEM, Gibco 10370-021) supplemented with 10% Fetal Bovine Serum (FBS, Atlantic Biological S12450), 1% Glutamax (Gibco 35050-061), 1% Pen Strep (Gibco 15140-122). Differentiation Media: MEM alpha (Gibco 12571-063) supplemented with 10% FBS, 1% Glutamax, 1% Pen Strep, 50ug/ul Ascorbic Acid (Sigma A4544-25G), 10mM beta-Glycerophosphate (Sigma G9422-100G), 10nM Dexamethasone (sigma D4902-25MG). RNA was isolated from ∼0.5×10(6) cells at days 0, 2, 4, 6, 8 and 10 of differentiation as recommended (RNAeasy Minikit. Qiagen 74106). Mineralized nodule formation was measured by staining cultures with Alizarin Red (40 mM, pH 5.6; Sigma A5533-25G). Reported results were obtained from three biological replicate experiments.

### RNA sequencing and Differential Gene Expression analysis

Computational analysis of RNA sequencing data for the 17 bone samples, Farr et al. ^24^ and the hFOB samples were performed using a custom bioinformatics pipeline. Briefly, FastqQC (http://www.bioinformatics.babraham.ac.uk/projects/fastqc/) and RSeQC ^40^ were used to assess the quality of raw reads. Adapter trimming was completed using Trimmomatic ^41^. Sequences were aligned to the GENCODE v34 ^42^ reference genome using the SNP and splice aware aligner HISAT2 ^43^. Genome assembly and abundances in transcripts per million (TPM) were quantified using StringTie ^44^. Differential expression analysis for the hFOB differentiation experiment was performed using DEseq2 ^45^ across all six differentiation time points using analysis of deviance (ANODEV) which is conceptually similar to analysis of variance (ANOVA). Differential expression analysis for the comparison between this study’s samples and the samples in the literature was performed using DEseq2 ^45^ standard approach.

### lncRNA discovery

The Coding Potential Assessment Tool (CPAT) ^27^ was used to assess the protein-coding potential of the novel transcripts assembled. In short, CPAT is a machine learning algorithm trained on a set of known human lncRNAs to identify novel putative lncRNAs based on shared sequence features. We used all known lncRNAs in the latest human genome assembly (GRCh38) as the training set. Novel transcripts with coding probability < 0.367 are regarded as lncRNAs in accordance with software authors. Novel lncRNAs with TPM < 1 were regarded as noise and discarded.

### Allelic Imbalance analysis

Reads were aligned to the GENCODE v34 ^42^ reference genome using the SNP and splice aware aligner HISAT2 ^43^. The resultant BAM files were then used as input for variant calling using the GATK pipeline ^46^. Briefly, duplicate reads were identified using MarkDuplicates. Next, reads spanning introns were reformatted using SplitNCigarReads to match the DNA aligner conventions. Then base quality recalibration was performed to detect and correct for patterns of systematic errors in the base quality scores. Finally, the variant calling and filtration step was performed using HaplotypeCaller. The resultant vcf file included only known and novel snps and reference bias was corrected using WASP ^47^. Briefly, mapped reads that overlap SNPs are identified. For each read that overlaps a SNP, its genotype is swapped with that of the other allele and it is re-mapped. If a re-mapped read fails to map to exactly the same location, it is discarded. The resultant corrected BAM and filtered VCF files were used as input for GATK ASEReadCounter to provide a table of filtered base counts at heterozygous sites for allele specific expression. Bases with a read depth less than 20 were discarded. In order to determine significance, a binomial test was performed and only heterozygous sites with FDR corrected p-value of <0.05 were considered significant.

### Transcriptome Wide Association Studies

We conducted a transcriptome-wide association study by integrating genome-wide SNP-level association summary statistics from a bone mineral density GWAS ^9^ with GTEx version 8 gene expression QTL data from 49 tissue types. We used the S-MultiXcan ^29^ approach for this analysis, to correlate gene expression across tissues to increase power and identify candidate susceptibility genes. Gene-level associations were identified at FDR correction < 0.05 and were further filtered using fastENLOC (described in for a regional colocalization probability > 0.1 in at least one tissue type.

### Bayesian colocalization analysis

We used fastENLOC, a faster implementation of ENLOC ^30^ to perform Bayesian colocalization analysis. We integrated summary statistics from the most recent (and largest) eBMD GWAS ^9^ and eQTL data from 49 GTEx tissues ^48^. We used the recommended regional colocalization probability (RCP) threshold of >0.1 as indication of significant overlap between SNP and eQTL.

## Notes

### Competing Interest Statement

The authors have declared no competing interest.

